# Exaggerated CpH Methylation in the Autism-Affected Brain

**DOI:** 10.1101/069120

**Authors:** Shannon E. Ellis, Simone Gupta, Anna Moes, Andrew B. West, Dan E. Arking

**Affiliations:** McKusick-Nathans Institute of Genetic Medicine, Johns Hopkins University School of Medicine, Baltimore, Maryland 21205, USA; Department of Neurology, University of Alabama at Birmingham, Birmingham, Alabama 35294, USA

## Abstract

The etiology of autism, a complex neurodevelopmental disorder, remains largely unexplained. Here, we explore the role of CpG and CpH (H=A, C, or T) methylation within autism-affected cortical brain tissue. While no individual site of methylation was significantly associated with autism after multi-test correction, methylated CpH di-nucleotides were markedly enriched in autism-affected brains (~2-fold enrichment at p <0.05 cut-off, p=0.002). These results further implicate epigenetic alterations in pathobiological mechanisms that underlie autism.

## Main Text

Autism is a heritable neurodevelopmental disorder affecting one in 68 individuals in the United States^1^. Recent genetic studies have identified a handful of genes that contribute to autism^2^ and gene expression studies have begun to unravel how altered gene expression manifests within the autistic brain^3,4^; however, the majority of risk remains unexplained. In addition to genetic causes, epigenetic mechanisms have been proposed to play an important role in the development of the disorder. Three lines of evidence initially supported this hypothesis. First, direct alterations in epigenetic pathways can dramatically alter early embryonic and neonatal neurodevelopment in the same critical periods as autism-associated changes in the brain^5^. Second, mutations in indirect epigenetic effectors can result in autism-spectrum and related disorders, such as Rett syndrome, Fragile X syndrome, and Angelman syndrome^6^. Finally, deficiencies in DNA methylation (DNAm), historically studied in CpG islands in gene promoters as an indicator of transcriptional repression, have previously been implicated in autism^7–9^.

Initial studies of methylation in autism were limited by the number of sites investigated, a lack of dynamic range in microarrays, the number of samples available for study, and the use of DNA that was procured from cell lines and tissue other than the brain. To gain a more complete picture of altered DNAm in autism, we carried out Reduced Representation Bisulfite Sequencing (RRBS) in 71 post-mortem cortical brain samples (BA19) at single nucleotide resolution with a quantitative measurement of DNAm across CpG-dense regions of the genome^10^. RRBS, in addition to querying methylation at more sites than the previously-used Infinium HumanMethylation450 array (Illumina)^11,12^, enables measurement of methylation at cytosines outside of the classically studied CpG context. While CpH methylation (mCH, where H=A,C, or T) is rare in most tissues, it accumulates in DNA in human and mouse brain postnatally, ultimately reaching levels similar to that of CpG methylation (mCG) in brain DNA^13–15^. In contrast to mCG, which remains largely unchanged during postnatal development, mCH accumulation correlates with synaptogenesis and increases especially during the first few years of life^13,14^, a time period of particular interest in autism. Thus, we used post-mortem cortical brains samples to characterize CpG and CpH methylation in autism-affected brain tissue and compared this to matched neurologically normal control brain tissue.

After the removal of sample outliers, 63 samples were included for analysis, comprising 29 autism cases and 34 controls (**Supplementary Table 1**). Methylation was estimated at cytosines with greater than 10 reads across at least 20 cases and 20 controls, yielding methylation estimates at 1.0M CpG and 3.3M CpH sites (**Supplementary Fig. 1**). No individual CpG or CpH sites were significantly differentially methylated after correction for multiple testing (**Supplementary Table 2-3**, **Supplementary Fig. 2**).

In addition to testing for differential methylation at individual sites, we measured global changes associated with hypo- or hypermethylation. Among sites demonstrating nominal differential methylation (p<0.05), there is a consistent and statistically significant proportion of cytosines demonstrating increased methylation within the CpH context (Fig. 1b, p=0.002 with 65.2% of sites demonstrating hypermethylation), but not the CpG context (Fig. 1a). Further, given that more stringent p-value cut-offs for differentially methylated sites should enrich for true positives, we hypothesized that the global hypermethylation signal would increase in strength with increasingly stringent p-value cut-offs in the CpH analyses, but not in the CpG analyses, which did not yield global differences. Indeed, as more stringent differential methylation p-value cutoffs were imposed, a greater skewing in the number of hypermethylated to hypomethylated sites was observed (Fig. 1b). As expected, this trend was not seen in the CpG sites (Fig. 1a). Moreover, the effect size of this hypermethylation signal increased with larger methylation differences between cases and controls (**Supplementary Fig. 3**). Taken together, these data suggest that small increases (CpH sites with a differentially methylated p-value < 0.001 demonstrate a median 1.8% increase in cases relative to controls) in methylation across many individual sites are found at cytosines outside of the classically studied CpG context in the autistic brain.

**Figure 1.**
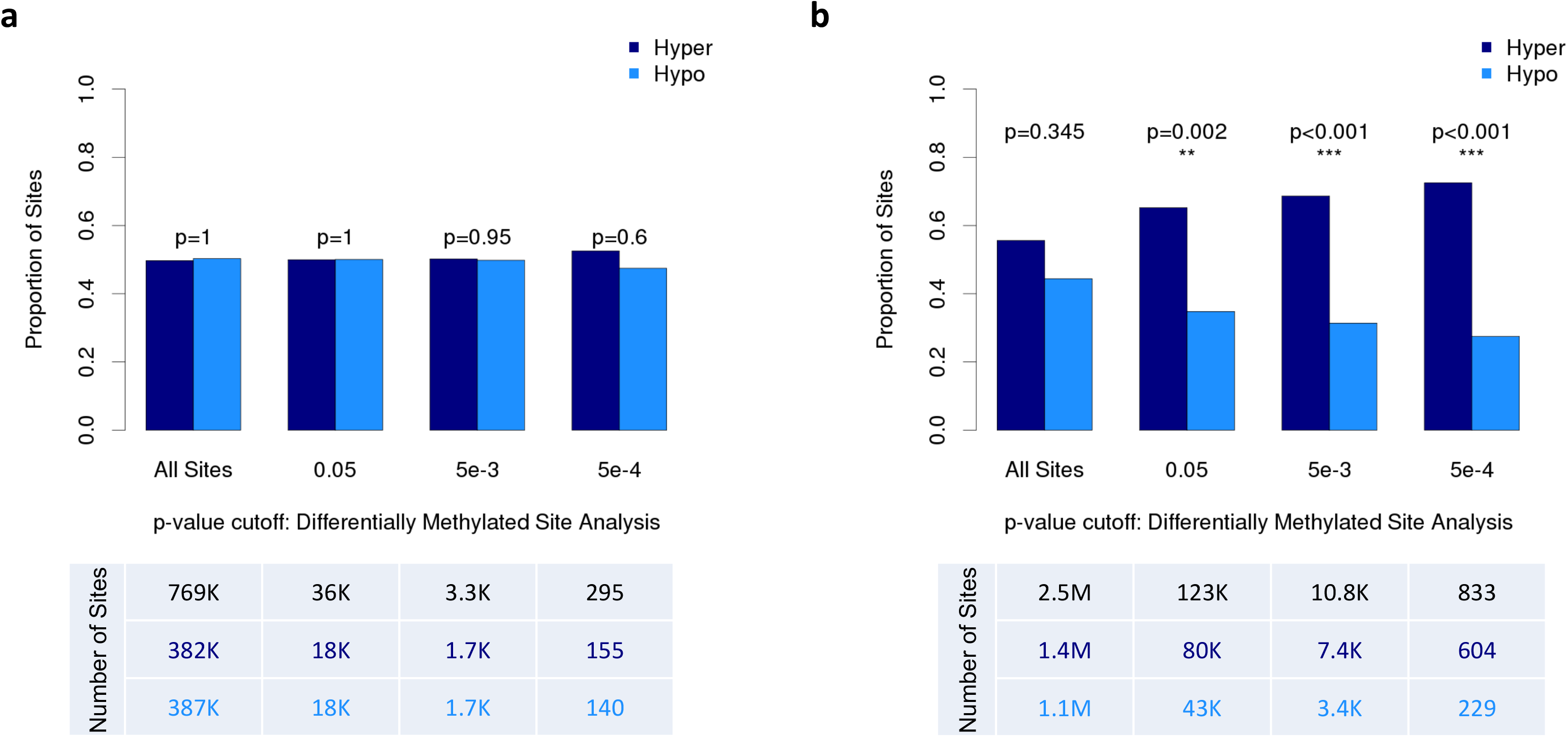
Proportion of hyper- and hypo-methylated sites in the CpG and CpH contexts. Proportion of sites (y-axis) across increasingly stringent differentially methylated p-value cutoffs (x-axis). The number of cytosines at each differentially methylated p-value cutoff are displayed in the tables (below). (**a**) With approximately half of all sites demonstrating increased methylation (navy) and the other half decreased methylation (light blue), CpG sites behave as expected under the null. This pattern holds across increasingly stringent differential methylation p-value cut-offs demonstrating no global differences in methylation within the CpG context. (**b**) The proportion of cytosines demonstrating hypermethylation is not significantly different from the proportion demonstrating hypomethylation when looking at all CpH sites; however, with increasingly stringent differentially methylated p-value cutoffs, there is a significant proportion of hypermethylated CpH sites in the autistic brain.

To better understand how altered mCH may be linked to the pathobiology of autism and aberrant neurodevelopment, we tested for enrichment of hypermethylated CpHs in various functional categories annotated across the genome. We used a Fisher’s exact test to detect enrichment of hypermethylated cytosines in 20 functional categories of the genome at several thresholds produced in the differential methylation analysis. This analysis highlights a role for increased methylation at CpH sites within repetitive regions of the genome (OR=1.39, p=5.7×10^−4^), in regions that contain non-polymorphic human-specific CpGs, termed beacons^16^ (OR= 1.27, p=0.04), and at deactivating histone marks in the brain (H3K27me3: OR=1.22, p=6.8×10^−3^; H3K9me3, OR=1.22, 1.6×10^−2^) (Fig. 2). Of note, histone-specific enrichment was not seen in any of the ten histone marks tested using data generated from a lymphoblastoid cell line, suggesting that this enrichment is tissue-dependent (**Supplementary Fig. 4**). Given previous reports of altered gene expression at transcriptional regulators^17^, these results not only corroborate previous findings but also further suggest a role for general transcriptional suppression at the level of mCH within the autistic brain. Further, as autism is a disorder that includes deficits in language, a key trait unique to humans, enrichment for mCH within regions known to harbor human-specific CpGs and at repetitive regions, which account for a substantial amount of variation between humans and other species, offers a particularly interesting avenue for further study. Taken together, this finding implicates increased methylation within autism brain tissue at cytosines outside of the canonical CpG di-nucleotide. It is not clear whether increased CpH methylation in autism is causal, protective, or benign in the etiology of disease. Given that mCH is specifically enriched in both the human and mouse brain^13^, future studies can begin to probe the function of CpH methylation in successful and aberrant neurodevelopment.

**Figure 2.**
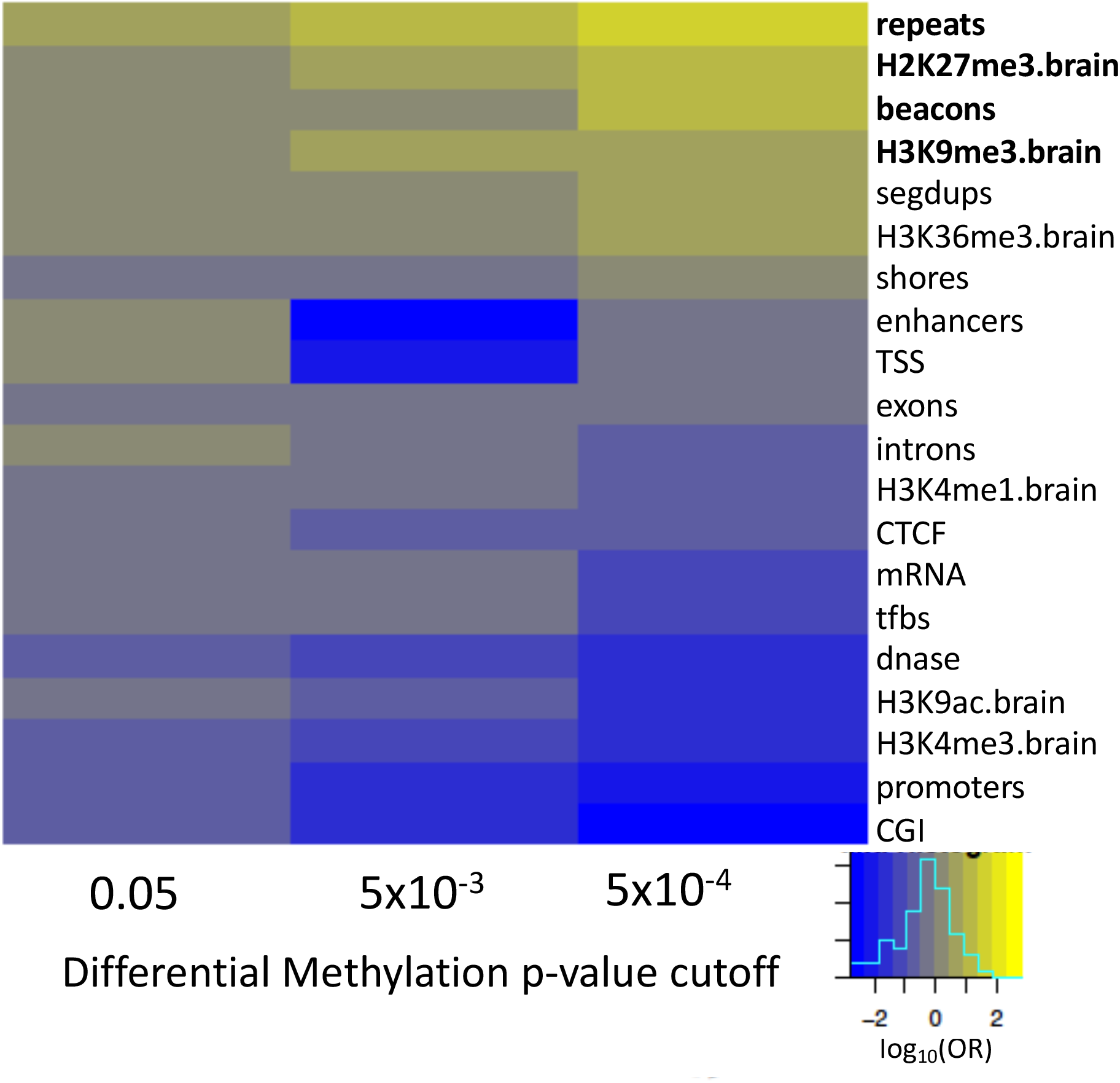
Genomic enrichment of hypermethylated CpH sites. For each genomic category, effect of enrichment (log odds ratio) is plotted across increasingly stringent differential methylation analysis p-value cutoffs (x-axis). Enrichment within a genomic category is indicated with the color yellow. Categories demonstrating significant enrichment (p<0.05) are in bold.

To maximize the number of samples that could be sequenced, this study employed RRBS rather than whole genome bisulfite sequencing (WGBS). As RRBS enriches for CpG rich regions of the genome, we are unable to estimate methylation for cytosines outside of CpG rich regions. As sequencing costs continue to decline, WGBS of all the available brain tissue specimens will become more feasible and will undoubtedly add further insight into the role of methylation and other epigenetic phenomenon in autism. Additionally, given the scarcity of samples, sample size is always a cause for concern in post-mortem brain studies. Here, we report findings from the largest number of samples studied to date. As such, we are 80% powered to detect mean methylation differences greater than or equal to 2.6% (**Supplementary Fig. 5**); however, group differences of smaller effect or idiosyncratic changes could have been missed in these analyses.

In summary, we report that increased CpH methylation occurs throughout the genome in DNA from autism-affected brain. These CpH sites are strongly associated with repetitive regions, deactivating histone marks, and beacons, offering new insights into how the epigenome may be affected in autism.

## Online Methods

### Samples

Samples were acquired through the Autism Tissue Program (which has since joined with the Autism Brain Net, https://autismbrainnet.org/). Post-mortem, frozen brain samples from the cerebral cortex Brodmann area (BA) 19 were collected at two different brain banks: the Harvard Brain Tissue Resource Center and the NICHD Brain and Tissue Bank at the University of Maryland with written informed consent having been obtained from next-of-kin or a legal guardian. Work herein was both approved by the IRB of The Johns Hopkins Hospital and University of Alabama at Birmingham and conducted in accordance with institutional guidelines.

### RRBS Library Preparation

Seventy-one samples were prepared for reduced representation bisulfite sequencing (RRBS). RRBS libraries were prepared using 100 ng of genomic DNA (gDNA). gDNA was first digested with MspI making cuts exclusively at methylated cytosines. 3’ A-overhangs were created and filled in with Klenow Fragments. DNA was then purified using the Qiagen MinElute Kit. Methylated ilAdap PE adapters (Illumina) were ligated to purified gDNA. Fragment size selection (105-185bp) was carried out by gel extraction on a 2.5% NuSieve GTG agarose gel (Lonza). DNA was purified using Qiaquick Gel extraction Kit eluting DNA in elution bugger pre-warmed to 55 degrees Celsius. Bisulfite treatment was performed using the ZymoResearch EZ DNA Methylation Gold Kit following manufacturer’s instructions; however, we eluted with 20μl M-Elution buffer. Bisulfite-treated DNA was cleaned up using EpiTect spin columns Samples were PCR amplified (using the following primers:

AATGATACGGCGACCACCGAGATCTACACTCTTTCCCTACACGACGCTCTTCCGATC

^*^T and

CAAGCAGAAGACGGCATACGAGATCGGTCTCGGCATTCCTGCTGAACCGCTCTTCCG

ATC^*^T; ^*^=phosphorothioate bond) and size selection was carried out on a 3% Metaphor Agarose Gel to ensure that fragments of the correct size (175-275bp) were amplified. PCR product was cleaned up using the Qiagen minElute column, eluting with elution buffer warmed to 55 degrees Celsius. Each sample (10nM) was sequenced in a single lane on the Illumina HiSeq2000 to produce 50bp single end reads.

### Alignment

Adaptor sequences were removed and reads shorter than 20 bp were excluded using Trim Galore! (v0.2.8). Remaining reads were aligned using Bismark (v0.7.7)^18^ allowing for one mismatch and setting the seed substring length to 24.

### Methylation Estimation

Two separate analyses were carried out based on cytosine context; one for cytosines in the CpG context and a separate analysis for all other cytosines in the genome (CpH). Samfiles for every sample and each of the two contexts were formatted for input into the R package ‘methylKit’^19^ (v0.9.5) using in-house scripts. Reads were filtered in methylKit based on read count discarding bases with coverage below 10X as well as those with coverage greater than the 99.9^th^ percentile of coverage in each sample to remove reads suffering from PCR bias. Data were normalized based on median coverage and methylation percentage estimated using ‘normalizeCoverage’ and ‘percMethylation’, respectively within methylKit.

### Illumina 27K Methylation Array

To independently verify methylation estimates from RRBS, CpG methylation was also analyzed in 71 cortical brain samples using the HumanMethylation27 BeadChip. These samples comprised 41 controls and 30 autism cases. Data were generated as described previously^20^. Normalized β–values were used for analysis. For comparison to RRBS data, mean methylation was quantified for the 1,249 CpGs that directly overlapping between the two platforms. Sites directly measured by both platforms had highly correlated measures of mean methylation (R^2^=0.92), offering confidence in the measurements acquired by RRBS (**Supplementary Fig. 6**).

### Sample Outlier Removal

Four samples were excluded from analysis upon initial diagnostics as their profiles indicated failed library preparation or failed sequencing. Two were removed due to the fact that nearly all (>99%) of their cytosines were methylated after alignment and methylation estimation (**Supplementary Fig. 7b-c**). A third sample was removed because its CpG methylation percentage distribution was not bimodal (**Supplementary Fig. 7d**). The fourth sample was removed because its read coverage distribution did not match the expected distribution (**Supplementary Fig. 7e**).

After identifying samples that failed library preparation and/or sequencing, remaining sample outliers were identified using surrogate variable analysis (SVA).^21^ Ten surrogate variables (SVs) were generated using methylation estimates from CpG sites with data across all samples (254,824 CpGs). Samples greater than four standard deviations away from the mean in any of the SVs generated were identified as sample outliers. This process was carried out iteratively, and after each round of sample outlier removal, the percentage of known brain meQTLs^22,23^ detected was quantified using a method previously developed for RNA-Sequencing data^24^ to guide data analysis. After each round of sample outlier removal, *cis* meQTLs (1Mb) were detected at SNPs and CpGs present in both the previously reported meQTL studies and the brain data using high quality genotype data described previously for these samples^24^. meQTLs were detected using MatrixEQTL^25^ with age, sex, site and SVs included as covariates, and the percentage of known meQTLs was recorded. This process enabled us to confidently move forward with 63 samples, including 29 autistic cases and 34 controls, as this sample size maximized the percentage of known meQTLs detected, in all downstream analyses (**Supplementary Fig. 8**).

### Single Site Differential Methylation Analysis

Methylation outliers at each single site were defined as any sample greater than three standard deviations away from the mean methylation at that site and removed. Only variant sites were included for analysis, excluding the 25% least variable sites from analysis. Single site differential methylation was then carried out on each site using the ‘*lmFit*’ function in the ‘limma’ R package^26^. For all cytosines, case-control status was regressed on methylation percentage with age, sex, brain bank, and ten SVs included as covariates (‘full model’). Ten SVs were generated using methylation data from all variant sites with data across all samples utilizing the “irw” method and protecting case-control status. Additionally, as read coverage impacts our confidence in methylation estimates, the log10 of read coverage at each site was included as weights in the model.

Statistical significance was determined by residual bootstrapping, again using ‘limma’. For each bootstrap, the full model (described above) was fit and residuals recorded. A null model, in which the variable of interest (here, case-control status) was excluded, was also fit. The residuals from the full model were resampled with replacement, randomizing the sample order. ‘Pseudonull’ data were then generating adjusting the fits from the null model with the resampled residuals from the full model. These pseudonull methylation values were then substituted as the outcome variable into the full model, generating a null set of p-values. These p-values were collected for each of the 1,000 bootstraps to empirically determine study-wide significance.

### Global Altered Methylation Analysis

For each cytosine context, the proportion of sites hypermethylated (defined as mean methylation in cases greater than zero) was calculated at three p-value cutoffs (0.05, 5×10^−3^, and 5×10^−4^). To assign significance, this proportion was then compared to the proportion of sites hypermethylated in each of the 1,000 residual bootstraps (**Supplementary Fig. 9**).

### Lists of Functional Genomic Categories

Lists for twenty-eight different functional genomic categories to test for enrichment of hypermethylated cytosines within the CpH context were downloaded from four different sources: (**1**) the UCSC Genome Browser (mRNA, transcription factor binding sites (tfbs), DNase I hypersensitive sites (dnase), enhancers, CTCF binding sites (CTCF), segmental duplications (segdups), repetitive regions (repeats), and histone marks from lymphoblastoid cell line GM12878 (H3K4m1, H3K4m2, H3K4m3, H3K9Ac, H3K9m3, H3K27Ac, H3K27m3, H3K36m3, H3K79m2, and H4K20m1), (**2**) UCL Cancer Institute (beacons), (**3**) the ‘methylKit’ package^19^ (promoters, exons, introns, transcription start sites (TSS), CpG Islands (CGI), and CGI shores) and (**4**) the Epigenome Roadmap Project^27^ (H327me3.brain, H3K9me3.brain, H3K36me3.brain, H3K4me1.brain, H3K9ac.brain, and H3K4me3.brain) (**Supplementary Table 4**). Brain data from the Epigenome Roadmap project were downloaded from adult cingulate gyrus. For histone marks with data generated on more than one individual (H3K36me3.brain, H3K4me1.brain, H3K4me3.brain, and H3K9me3.brain), the intersection of regions across individuals was utilized for downstream analyses.

### Functional Enrichment Testing

To test for genomic enrichment of hypermethylated CpH sites in each genomic list and at each p-value cutoff from the differential methylation analysis (0.05, 5×10^−3^, and 5×10^−4^), a two-sided Fisher’s exact 2×2 test was carried out. For each list and at each differential methylation p-value cutoff, odds ratios and p-values for enrichment were recorded.

### Power Calculation

Power calculations were carried out using the “pwr.t2n.test” function from the ‘pwr’ package in R. This two-sided t-test of means for samples of different sizes (N=34 controls and 29 cases) was carried out at the 0.05 significance level (Type I error probability).

